# CHD1 is a synthetic lethal vulnerability in MYC-driven breast cancer

**DOI:** 10.1101/2025.08.14.670376

**Authors:** Brandon Cho, Giacomo Furlan, Peter Lin, Mélissa Shen, Kirti Mittal, Andrew Mazzanti, Linda Z. Penn, Miguel Ramalho-Santos

## Abstract

The MYC transcription factor is a key regulator of growth during development and a potent cancer driver when its expression is dysregulated. Strategies to inhibit MYC oncogenic activity would mark a significant advance, but decades of efforts to target MYC directly have not been fruitful. Understanding how MYC drives transformation and tumor growth may provide new therapeutic avenues in a variety of cancers. By intersecting two independent genome-wide screens, we identified loss of the chromatin remodeler Chromodomain-Helicase DNA-binding 1 (CHD1) as a potential synthetic lethal target in MYC-driven breast cancer. Knockdown of *CHD1* in a xenograft model of MYC-driven breast cancer suppresses tumor growth in vivo. In tissue culture models, we found that knockdown of *CHD1* suppresses cell proliferation and induces cell death, specifically when *MYC* is overexpressed. Mechanistically, we found that CHD1 is required to maintain an open chromatin landscape and a transcriptional program associated with cancer progression in *MYC* overexpressing breast cells. Follow-up experiments indicate that this synthetic lethality may arise from nucleolar stress and p53 activation. These findings provide new insights on the chromatin-level regulation of MYC-driven breast cancer and uncover CHD1 as a novel synthetic vulnerability and potential therapeutic target.

## INTRODUCTION

The c-MYC (MYC) is a transcription factor is a master regulator that plays essential roles in cell growth, proliferation, apoptosis and metabolism during embryonic development and cell differentiation^1–6^. Given this wide range of functions, the expression and activity of MYC is under tight regulation. When this regulation is defective and MYC is overexpressed, it becomes a potent oncogene that drives transformation in a wide variety of cancers^7,8^. This dysregulation of MYC can occur through various mechanisms, including direct *MYC* gene amplification (e.g., via extrachromosomal DNA) and activation (e.g. Ras) or loss (e.g. PTEN) of upstream signaling cascades. MYC dysregulation occurs in the majority of human cancers, including breast, colon, cervix, lung and pancreatic cancer, as well as in several types of leukemias and lymphomas^9^. However, efforts to develop inhibitors to target this potent oncoprotein have been hindered by its intrinsically disordered structure and the lack of enzymatic pockets. Thus, despite decades of research, drugs that directly inhibit MYC have yet to be approved for patient treatment^10,11^.

An alternative route to targeting MYC-driven cancer is to identify synthetic lethalities, whereby a particular perturbation hinders cancer progression specifically when MYC is overactivated, but not in normal cells^12^. Several genome-wide screens have been conducted to identify genes whose loss is synthetic lethal with MYC activation^12–18^. Using this strategy, factors such as SUMO-activating enzyme (SAE1/2), YTH N6-methyladenosine RNA binding protein 2 (YTHDF2), and Topoisomerase I (TOP1) have been shown to be functionally required in various MYC-driven cancer models^17–20^. Thus, identifying genes that are synthetic lethal with MYC dysregulation can reveal unique vulnerabilities of MYC-driven cancers that may be exploited to development novel therapeutics.

In this study, we identified Chromodomain Helicase DNA-binding protein 1 (CHD1) as a potential vulnerability of MYC-driven breast cancer. CHD1 is a chromatin remodeler associated with active gene promoters that maintains open chromatin and facilitates transcriptional output by both RNA Pol I and II^21–24^. CHD1 is not required for basal transcription, but plays essential roles in developmental transitions associated with rapid growth and high transcriptional output, or hypertranscription^21–28^. Here, we report that CHD1 is required for MYC-driven breast epithelial cell transformation in vitro and in vivo. Our data further reveal that CHD1 is required to maintain a chromatin and transcriptional state associated with cancer progression and invasion. These findings indicate that CHD1 is a novel candidate therapeutic target in MYC-driven cancer.

## RESULTS

### *CHD1* knockdown suppresses growth of MYC-driven breast tumors in vivo

We compared two independent genome-wide synthetic lethal screens in MYC-driven breast cancer models, one using RNA interference and another using CRISPR knockout^13,20^. Nineteen genes were commonly identified as candidates in both screens, with *CHD1* being one of them (Fig. 1A, B). Specifically, *CHD1* was ranked third among these 19 shared genes, suggesting a possible synthetic lethality with *MYC* (Fig. 1B). Given the role of CHD1 in growth of stem and progenitor cells during development^22,23,27^, we set out to further explore the potential synthetic lethal interaction between *MYC* and *CHD1*. For this purpose, we used an established xenograft model of MYC-driven breast cancer^20,29^. This model employs MCF10A cells, which are non-transformed human basal breast epithelial cells, into which two of the most common mutations in breast cancer have been introduced: an activating mutation in the phosphoinositide 3 kinase pathway (PIK3CA^H1047R^), and a vector driving ectopic *MYC* expression^29^. Cells with both the <u>P</u>IK3CA^H1047R^ mutation and ectopic <u>M</u>YC, referred to as MCF10A.PM (PM), emulate primary human invasive ductal carcinoma at both the pathological and molecular levels. This oncogenic activity is driven and dependent on dysregulated MYC expression as cells expressing only the <u>P</u>IK3CA^H1047R^ mutation and <u>e</u>mpty vector, referred to as MCF10A.PE (PE), form normal breast acinar structures (Fig. 1C). One notable characteristic of this model is that the PM cells develop robust tumors upon xenograft into mice, while the PE cells do not^29^. We developed lentiviral vectors for doxycycline (Dox)-inducible short hairpin-RNA (shRNA) to knockdown (KD) *CHD1*. Cell lines carrying either of two independent shRNAs for *CHD1* KD (shCHD1 A and B) or a control shRNA targeting GFP (shGFP) were generated. KD efficiency was confirmed by both qRT-PCR and Western blotting (Fig. S1A-B).

**Figure 1.**
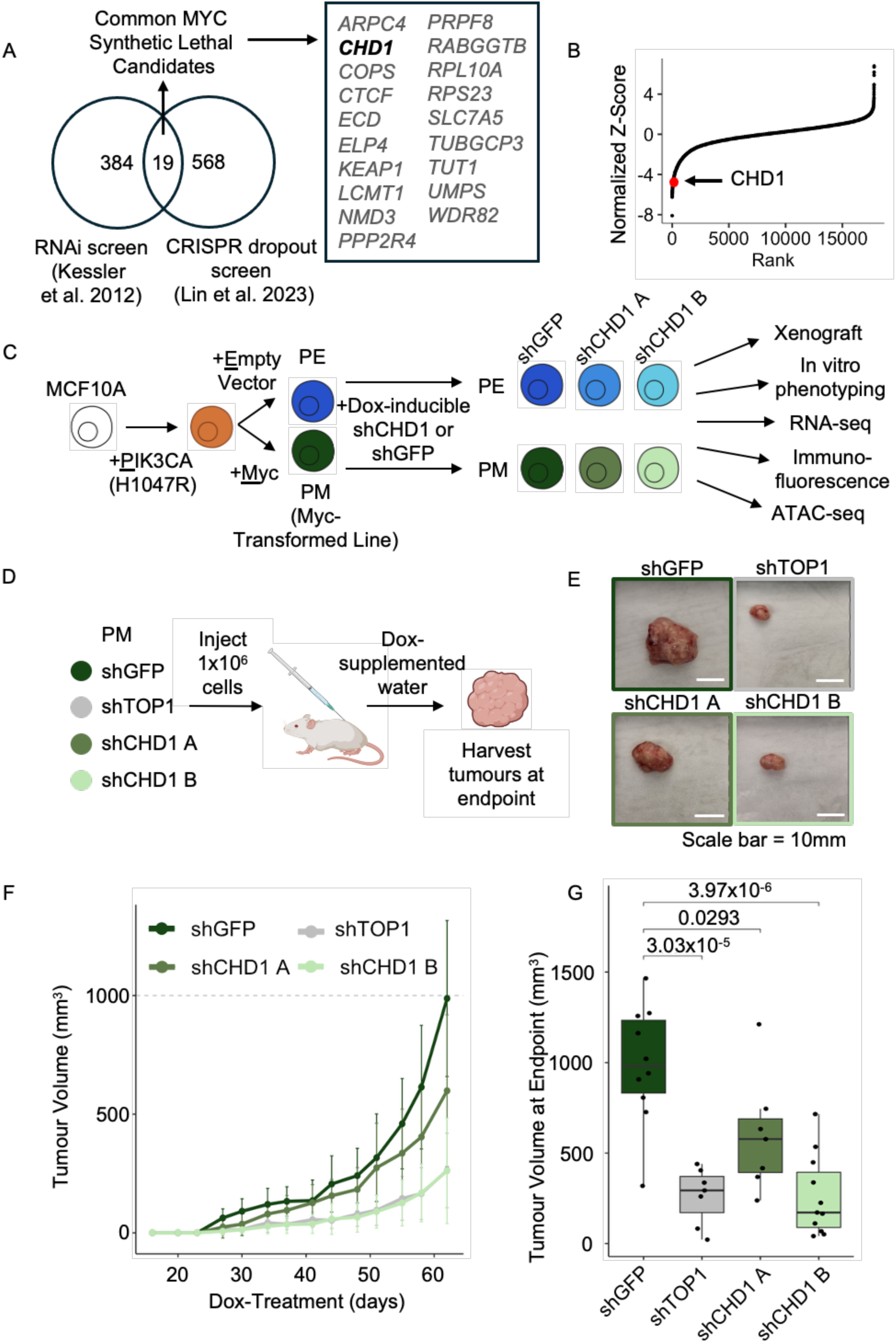
*CHD1* knockdown suppresses growth of MYC-driven breast tumors in vivo. A) Venn diagram overlapping the number of common synthetic lethal candidates in MYC-driven breast cancer models identified in two independent screens: an RNAi screen (Kessler et al. 2012) and a CRISPR dropout screen (Lin et al. 2023). CHD1 is one of 19 potential candidates identified between these two screens. B) CHD1 ranks 81 based on normalized Z-score from the Lin et al. 2023 CRISPR knockout screen, and third from the group of 19 genes shared between the two screens. C) Schematic of the MCF10A.PE/PM MYC-driven breast cancer model using a Dox-inducible shRNA knockdown system. Two independent *CHD1* KD lines were generated targeting different regions of the gene. shCHD1 A and B target exon 24 and 28, respectively. D) Schematic of xenograft experiments. A panel of 4 different conditions in the MYC-driven PM line were included. shGFP (negative control), shTOP1 (positive control), and two shCHD1 conditions (shCHD1 A and shCHD1 B). E) Representative images of excised tumors at endpoint, scale bars represent 10mm. F) Tumor volume (mm^3^) measured over the course of a xenograft experiment with humane endpoint determined to be at 62 days of Dox treatment, n=7-11 mice per condition. Error bars represent standard deviation. G) *CHD1* KD significantly reduces tumor volume at endpoint (mm^3^). P-values were calculated by one-way ANOVA with Tukey’s HSD.

A panel of validated control or *CHD1* KD PM cells were then xenografted into mice, with Dox supplemented in the drinking water (Fig. 1D). Topoisomerase 1 (*TOP1*) was recently identified as a MYC-synthetic lethal target in this model, and therefore an shTOP1 line was included as a positive control^20^. Interestingly, we found that both of the *CHD1* KD groups displayed a notable suppression of tumor growth and endpoint volume (Fig. 1E-G, Fig. S1C-D). This is particularly the case for shCHD1 B, which yielded results very similar to that observed in shTOP1. These data indicate that *CHD1* is a synthetic lethal vulnerability in MYC-driven breast cancer in vivo.

### *CHD1* knockdown is synthetic lethal with MYC-overexpression in breast cells in vitro

We next sought to explore further the genetic relationship between *MYC* and *CHD1* using cell culture models. *CHD1* KD significantly impaired the proliferation of PM cells, but not PE cells (Fig. 2A). This effect was recapitulated in clonogenic assays, where the two independent *CHD1* shRNAs reduced the colony-forming ability of PM cells, while having no impact on PE cells (Fig. 2B-C). This effect on colony formation was not seen when cells were not treated with Dox, validating the inducibility of the KD and the dependence of *CHD1* for MYC-driven cell proliferation (Fig. S2A). One feature of PM cells is that in 3D cultures they form disorganized acinar structures in vitro that are indicative of a transformed phenotype^29^. We found that *CHD1* depletion significantly attenuates the acinar-like transformation of PM cells in three-dimensional (3D) culture, whereby the cell aggregates instead maintained a smooth, spherical architecture upon *CHD1* KD (Fig. 2D-E, S2B).

**Figure 2.**
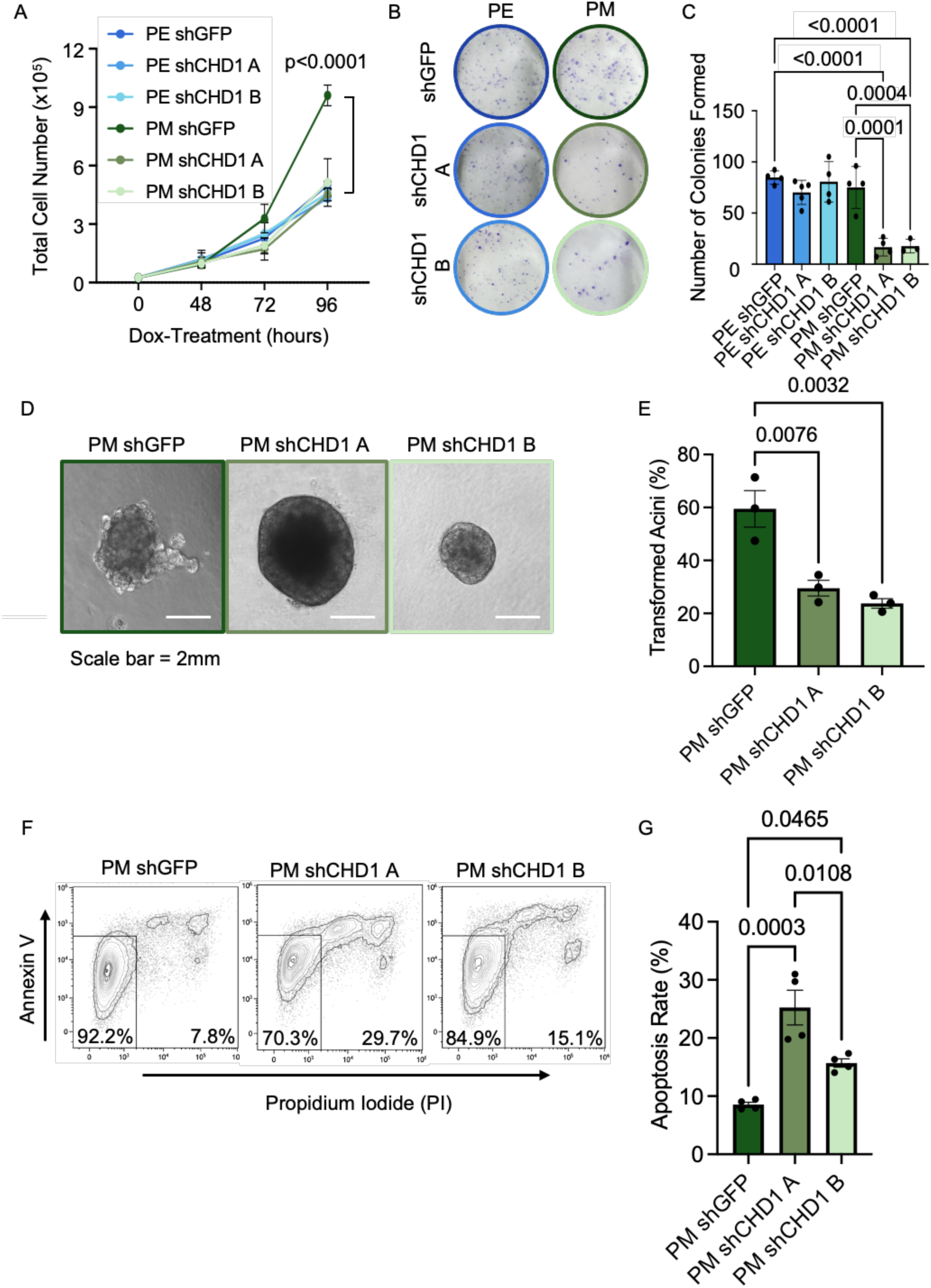
*CHD1* knockdown is synthetic lethal with MYC-overexpression in breast cells in vitro. A) Proliferation curve measuring total cell counts over a period of 96h of Dox-induced shRNA expression, n=3 independent biological replicates per timepoint. *CHD1* KD significantly impairs expansion of PM but not PE cells. P-values were calculated at 96h by one-way ANOVA with Tukey’s multiple comparisons test. B) Representative images of a clonogenic assay for the entire panel of PE and PM cells. C) Number of colonies formed in a clonogenic assay with cells treated with Dox, n=3-5 biological replicates per condition. *CHD1* KD significantly impairs colony formation ability of PM but not PE cells. Data are presented as mean ± SEM. P-values were calculated by one-way ANOVA with Tukey’s multiple comparisons test. D) Representative images of a 3D Matrigel growth assay of PM breast cancer cells cultured for 14 days. Scale bar represents 2mm. E) Quantification of 3D Matrigel growth assay in PM shGFP, PM shCHD1 A, and PM shCHD1 B cells after 14 days of tamoxifen treatment. Images were quantified based on the percentage of transformed acinar structures per condition. *CHD1* KD significantly reduces acinar transformation of PM cells. Data are presented as mean ± SEM. n=3 independent biological replicates. P-values were calculated by one-way ANOVA with Tukey’s multiple comparisons test. F) Example contour plots of flow cytometry data from AnnexinV/PI apoptosis assay. G) Cell death in the 10A.PM panel was measured by AnnexinV/PI flow cytometry at 96h of shRNA expression. Apoptosis rate was measured as all cells positive for PI, AnnexinV, or both. *CHD1* KD significantly induces apoptosis in PM cells. Data are presented as mean ± SEM. n=3 biological replicates. P-values were calculated by one-way ANOVA with Tukey’s multiple comparisons test.

To determine whether the defect in the ability of the PM cell lines to expand and transform upon *CHD1* KD is associated with cell death, we performed an AnnexinV/PI flow cytometry assay to assess apoptosis. The results reveal a significant increase in apoptosis rate upon Dox-induced *CHD1* KD in both PM shCHD1 A and PM shCHD1 B lines (Fig. 2F-G). Taken together, these data indicate that *CHD1* KD in PM cells significantly reduces their survival, growth and transformation under both two-dimensional and 3D growth conditions, further supporting the anti-tumor activity of *CHD1* KD in this MYC-driven breast cancer model.

### *CHD1* knockdown rewires the transcriptome of *MYC* overexpressing cells

We next determined the impact of *CHD1* KD in PE and PM cells at the transcriptomic level, by performing RNA-sequencing (RNA-seq). Principal component analysis and hierarchical clustering of the data revealed good correlations between replicates and independent shRNAs (Fig. 3A-B). We observed two clear large-scale transcriptional shifts: i) upon *MYC* overexpression (between PE and PM cells); and ii) upon *CHD1* KD (between shGFP and shCHD1 cells) (Fig. 3A-B). Given the synthetic lethality observed between *CHD1* KD and *MYC* dysregulation (Fig. 1A-B, 2A-C), we further explored the transcriptional changes induced by *CHD1* KD in PM cells (Fig. 3C, Supp. Table 1). Gene ontology and gene set enrichment analyses revealed that *CHD1* KD induces significant changes in transcriptional signatures of cancer hallmark pathways. These include upregulation of the p53 pathway and downregulation of cell cycle pathways such as E2F targets, G2M checkpoint and DNA replication, as well as TNFα pathway and inflammatory response (Fig. 3D-G, S3A-B). These data indicate that *CHD1* KD rewires the transcriptome of *MYC* overexpressing cells in ways consistent with the tumor-suppressive effect observed in xenografts (Fig. 1), with downregulation of the cell cycle and induction of p53.

**Figure 3.**
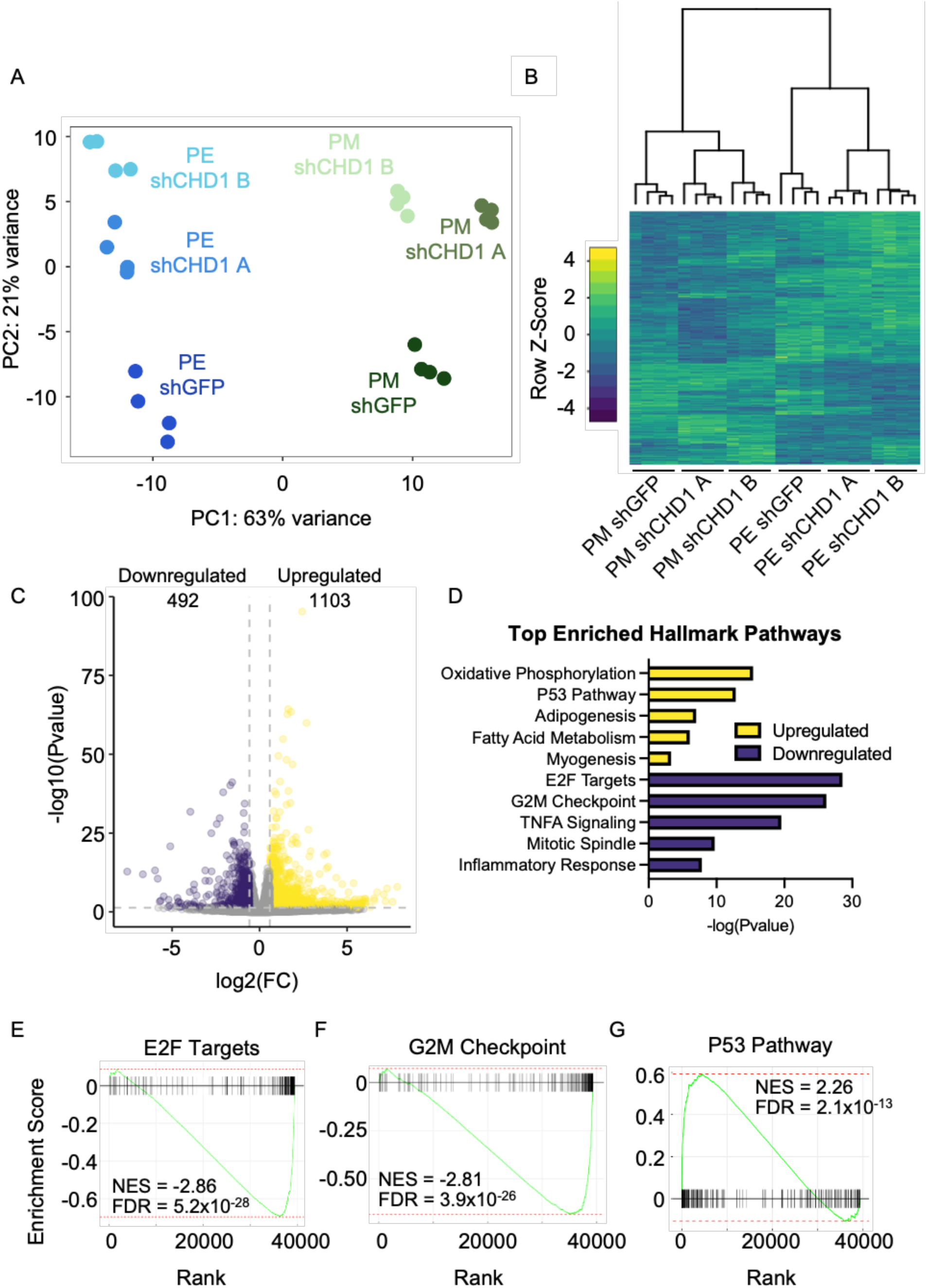
*CHD1* knockdown rewires the transcriptome of *MYC* overexpressing cells. A) Principal Component Analysis of RNA-seq data from 10A.PE and 10A.PM with either shGFP or shCHD1 induction for 72h. Each group has 4 biological replicates (n=4). Sample replicates are well correlated, *MYC* overexpression drives PC1 and *CHD1* KD drives PC2. B) Hierarchical clustering and heatmap of the top 2500 most variable genes in RNA-seq data from 10A.PE and 10A.PM with either shGFP or shCHD1 KD for 72h. Similarly to A), the two shRNAs targeting *CHD1* have concordant effects. C) Volcano plot of differential gene expression in the PM shCHD1 cell lines relative to PM shGFP line. There are 1103 significantly upregulated genes and 492 significantly downregulated genes. The horizontal dotted line indicates an FDR-corrected P-value of 0.05, while the vertical dotted lines indicate absolute fold changes of 1.5. D) Gene Ontology (GO) analysis of genes differentially expressed in the 10A.PM CHD1 KD conditions relative to the PM shGFP controls. Shown are the top 5 most upregulated (yellow) and downregulated (purple) Hallmark Pathways ranked by –log(Pvalue). E) Gene Set Enrichment Analysis (GSEA) of the Hallmark E2F Targets pathway in the 10A.PM CHD1 KD conditions relative to the PM shGFP control. F) GSEA of the Hallmark G2M Checkpoint pathway in the 10A.PM CHD1 KD conditions relative to the PM shGFP control. G) GSEA of the Hallmark P53 Pathway in the 10A.PM CHD1 KD conditions relative to the PM shGFP control.

### *CHD1* knockdown induces cell cycle defects and p53 expression

We followed up on the RNA-seq results by carrying out direct analyses of the cell cycle and p53 protein expression in *CHD1* KD cells. Using flow cytometry, we found that *CHD1* KD induces a significant reduction in the proportion of cells in S phase, relative to the PM shGFP controls, although this reduction in S phase is not attributable to specific increases in G1 or G2 cell cycle states (Fig. 4A-B, S4A). In parallel, immunofluorescence revealed a heterogeneous induction of p53 protein in PM *CHD1* KD cells relative to controls, in agreement with the RNA-seq data. This further supported by the increased levels of p21, a cell cycle inhibitor and key component of the p53 pathway, in *CHD1* KD cells (Fig. S4B). Together with the higher levels of apoptosis observed in *CHD1* KD cells (Fig. 2F), these data suggest that the growth suppressing effects of *CHD1* depletion in the context of *MYC* overexpression are mediated by a combination of p53 activation, cell cycle defects and increased cell death.

**Figure 4.**
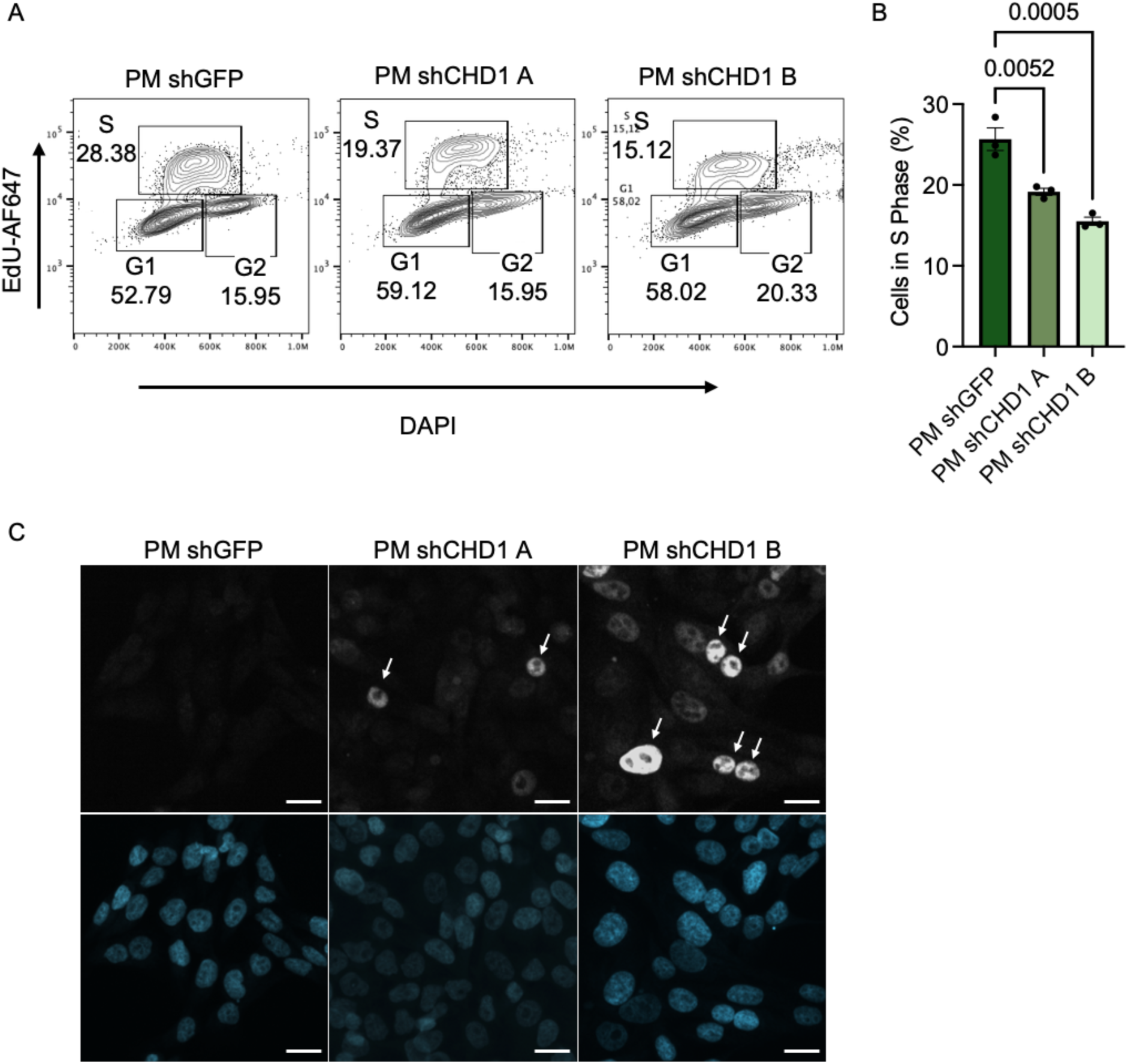
*CHD1* knockdown induces cell cycle defects and p53 expression. A) Representative examples of flow cytometry contour plots showing the gating strategy and data for cell cycle analysis. The x- and y-axes represent DAPI and EdU-AF647 fluorescence intensity, respectively. n=3 biological replicates. B) Quantification of the proportion of cells in S-phase from EdU cell cycle analysis by flow cytometry (n=3 biological replicates). *CHD1* KD leads to a significant reduction in the proportion of cells in S-phase. Data are presented as the mean ± SEM. P-values were calculated by one-way ANOVA with Tukey’s multiple comparisons test. C) Representative images showing induction of p53 (upper row) by *CHD1* KD in PM cells, as assessed by immunofluorescence and DAPI (lower row). Scale bars represent 20µm.

### The chromatin accessibility landscape of *MYC* overexpressing breast cells is dependent on *CHD1*

A core function of CHD1 is to remodel the chromatin landscape towards a more open, accessible state^21,24^ Therefore, we sought to explore the impact of *CHD1* KD on chromatin accessibility of PM cells by performing Assay for Transposase-Accessible Chromatin with sequencing (ATAC-seq, Fig. S5A-S5B). These results revealed that *CHD1* KD induces a significant loss in chromatin accessibility of PM cells, relative to controls (Fig. 5A). 880 open chromatin regions lost accessibility upon *CHD1* KD, whereas only 52 regions gained accessibility (Fig. 5B, Supp. Table 2).

**Figure 5.**
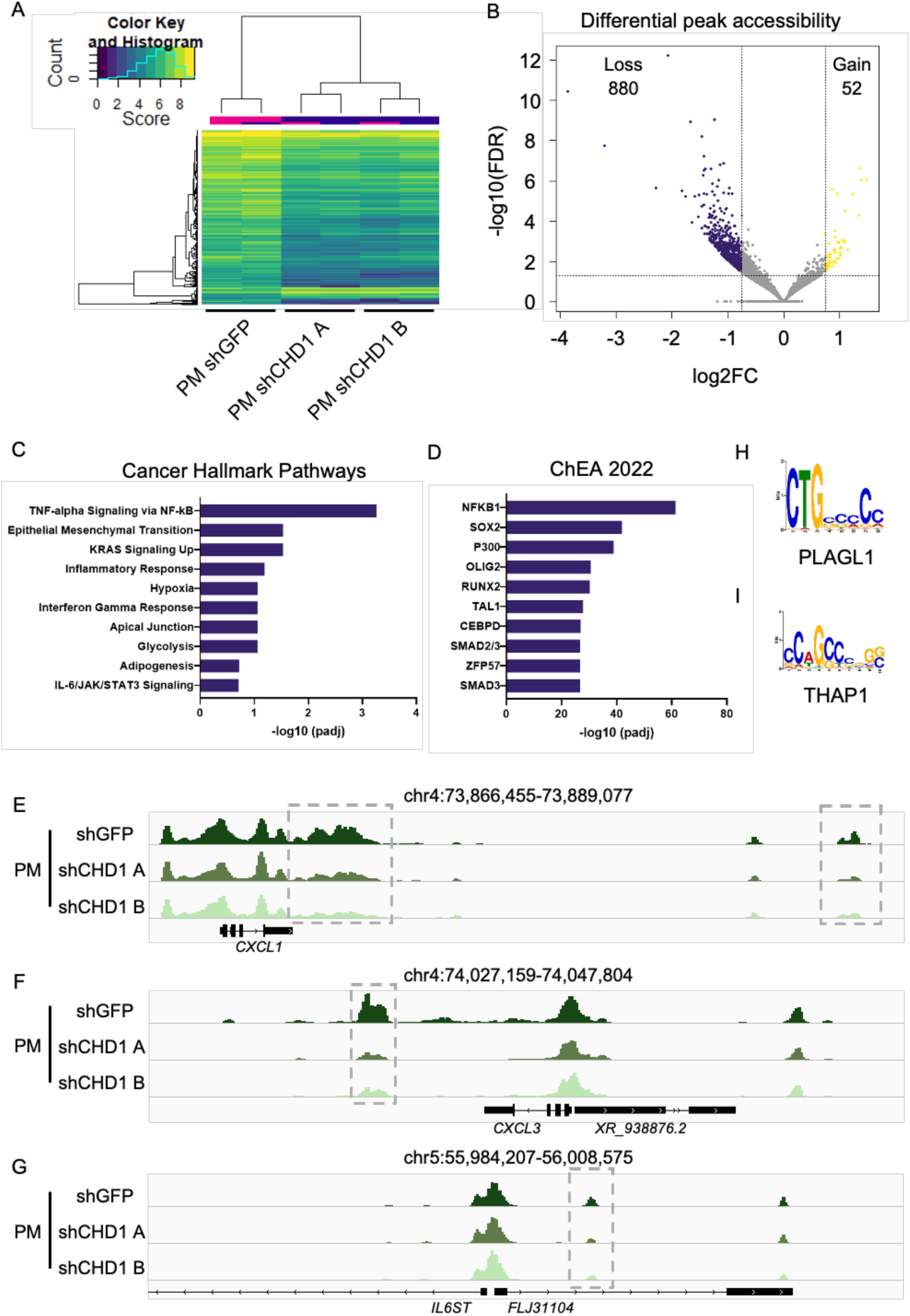
The chromatin accessibility landscape of *MYC* overexpressing breast cells is dependent on *CHD1*. A) Heatmap of the top 500 differentially accessible regions from ATAC-seq performed on PM shGFP, PM shCHD1 A and PM shCHD1 B cells after 72h of *CHD1* knockdown. The two shRNAs targeting *CHD1* have concordant effects and mostly induce loss of chromatin accessibility. Each group consists of 2 biological replicates. B) Volcano plot depicting differential ATAC-seq peaks between PM shCHD1 cells and PM shGFP cells. There are 880 regions with significant loss of chromatin accessibility and only 52 with gain. The horizontal dotted line indicates an FDR-corrected P-value of 0.05, while the vertical dotted lines indicate absolute fold changes of 0.75. C) GO analysis of regions with a significant loss of accessibility in the 10A.PM CHD1 KD conditions relative to the PM shGFP controls. Shown are the top 10 most significantly enriched Cancer Hallmark Pathways ranked by -log10(padj). D) ChIP enrichment analysis (ChEA 2022 dataset in Enrichr) of regions with a significant loss of chromatin accessibility upon *CHD1* KD relative to control cells, ranked by -log10(padj). E) Genome browser view of the region surrounding *CXCL1* on chromosome 4. Shown are merged data from the ATAC-seq replicates of shGFP control and shCHD1 lines. F) Genome browser view of the region surrounding *CXCL3* on chromosome 4. Shown are merged data from the ATAC-seq replicates of shGFP control and shCHD1 lines. G) Genome browser view of the region surrounding *IL6ST* on chromosome 5. Shown are merged data from the ATAC-seq replicates of shGFP control and shCHD1 lines. H) The top DNA motif that is most enriched in regions with a gain of chromatin accessibility upon *CHD1* KD with the transcription factor that best matches the motif: PLAGL1. I) The DNA motif that is the second most enriched in regions with a gain of chromatin accessibility upon *CHD1* KD with the transcription factor that best matches the motif: THAP1.

Given these findings, we first focused on regions of reduced chromatin accessibility following *CHD1* KD. These regions are enriched for genes of the inflammatory response, in agreement with the RNA-seq results (Fig. 3D, 5C). Regions of loss of chromatin accessibility are also associated with other cancer-associated pathways^30,31^, particularly those that been attributed to MYC activity such as the epithelial-mesenchymal transition, KRAS signaling and glycolysis, potentially contributing to the anti-tumor effects of *CHD1* KD. In agreement with these findings, regions with loss of chromatin accessibility are enriched for binding of transcription factors associated with inflammation (NFκB), stemness and oncogenic transformation (SOX2) and global transcription (P300), among others. Moreover, binding motifs for transcription factors that mediate inflammation (RELA, subunit of NFκB, and BATF) are enriched in ATAC-seq peaks that lose chromatin accessibility upon *CHD1* KD (Fig. 5C-D). Examples of genes of the inflammatory response located in the vicinity of regions that lose chromatin accessibility upon *CHD1* KD include *CXCL1* and *CXCL3* (Fig. 5E-F). *CXCL* genes are implicated in various cancers, including breast cancer, and are thought to promote cancer progression via activation of the TNFα pathway^32^. Similarly, *IL6ST,* which codes for the interleukin-6 receptor gp130 and is a prognostic biomarker of breast cancer biomarker^33,34^, also displays a region in its vicinity with decreased chromatin accessibility upon *CHD1* depletion (Fig. 5G).

To gain insights into the few regions that gain chromatin accessibility upon *CHD1* KD, we analyzed their potential enrichment for transcription factor binding motifs. The top two most enriched DNA-binding correspond to *PLAGL1* and *THAP1* (Fig. 5H-I). Interestingly, PLAGL1 is a zinc finger protein that interacts with p53 and p21^35^, both of which are increased in *CHD1* KD cells (Figs. 3D and G, 4C, S4C). THAP1 is a nuclear proapoptotic factor^36,37^, possibly paralleling our findings on increased apoptosis in *CHD1* KD cells (Fig. 2F). Taken together, these data indicate that CHD1 is required to maintain an open chromatin landscape in *MYC* overexpressing breast cells at genes belonging to pathways that drive cancer progression, most prominently the inflammatory response.

### Expression of *CHD1* is important for maintaining nucleolar function in MYC-driven breast cancer

We next sought to explore what aspects of nuclear biology may be reflected in the reprogramming of chromatin accessibility observed upon CHD1 depletion. We noticed that the signature of transcription factor binding sites enriched in regions that lose chromatin accessibility in *CHD1* KD cells (Fig. 5D), is nearly identical to that observed upon silencing of *LINE1* elements located in acrocentric chromosomes in human ESCs, which disrupts the nucleolar structure and function^38^. A further similarity between these two distinct contexts is that KD of *LINE1* in hESCs also leads to induction of p53, potentially as a consequence of nucleolar stress^38–40^. We therefore investigated the potential impact of *CHD1* KD on nucleolar function. Under nucleolar stress conditions, nucleoli marked by Nucleophosmin 1 (NPM1) form large, ring-like structures^41,42^. Immunofluorescence for NPM1 revealed a significant increase in such ring-like nucleoli in the PM shCHD1 cells, relative to control cells (Fig. 6B-C).

**Figure 6.**
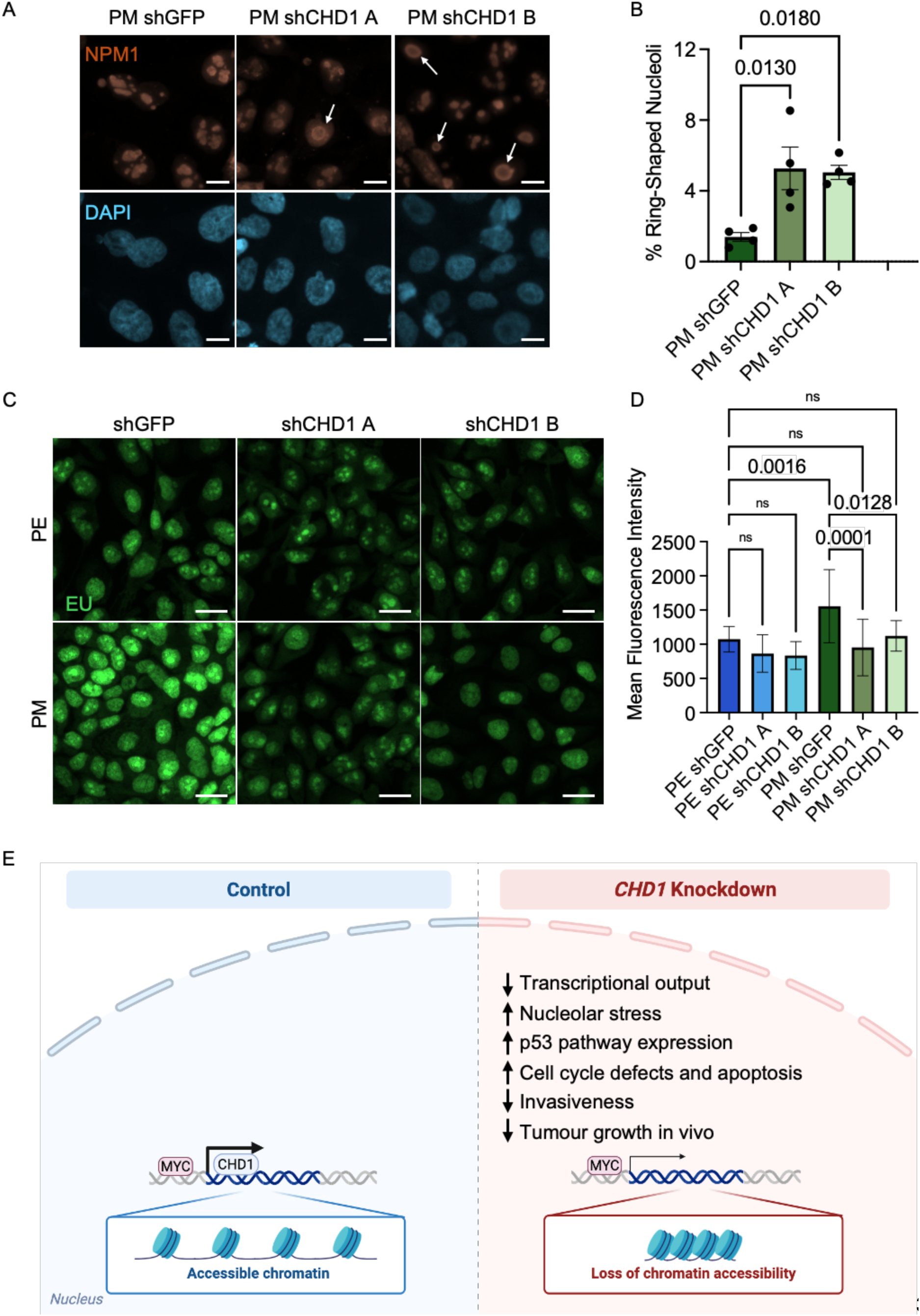
Expression of *CHD1* is important for maintaining nucleolar function in MYC-driven breast cancer. A) Representative images of NPM1 by immunofluorescence at 72h of Dox-treatment in control vs *CHD1* KD PM cells. White arrows depict “ring-shaped” nucleoli, indicative of nucleolar stress. Scale bars represent 10µm. B) *CHD1* KD induces a significant increase in the percentage of ring-shaped nucleoli. n=4 independent biological replicates. P-values were measured by one-way ANOVA with Tukey’s HSD. Data are presented as mean ± SEM. C) Representative images of immunofluorescence capturing EU incorporation, a measure of nascent transcription, in 10A.PE and 10A.PM cell lines at 72h of Dox-treatment. Scale bars represent 20 µm. n=3 independent biological replicates. D) Quantification of EU mean fluorescence intensity (MFI), n=3 independent biological replicates. MYC overexpression induces total nascent transcription, and this effect is suppressed in both shRNA CHD1 conditions. Data are presented as mean ± SEM. P-values were calculated by one-way ANOVA with Tukey’s multiple comparisons test. E) Model for the effect of *CHD1* KD in MYC-driven breast cancer. By analysing two unbiased genome-wide screens, we identify and validate the chromatin remodeler CHD1 as a synthetic lethal vulnerability in a xenograft model of MYC-driven breast cancer. Knockdown of CHD1 results in reductions in chromatin accessibility and nascent transcription, while inducing nucleolar stress, activation of the p53 pathway, cell cycle defects and apoptosis, ultimately suppressing invasiveness and tumor growth. Notably, the lethality induced by CHD1 KD is specific to breast epithelial cells overexpressing MYC. These results provide new insights into the chromatin-level regulation of MYC-driven breast cancer and uncover CHD1 as a novel candidate therapeutic target.

The nucleolus is the site of ribosomal RNA (rRNA) transcription by RNA Polymerase I, which accounts for the vast majority of nascent transcription in any given cell^43^. Moreover, global transcription, including of rRNA, is regulated by Chd1 in mouse ESCs^22,23^. We therefore assessed nascent transcription using 5-ethynyl uridine (EU) incorporation. Relative to PE cells, PM shGFP cells have significantly higher levels of nascent transcription (Fig. 6C-D), as would be expected for MYC overexpression^4,44^ Interestingly, this increase in nascent transcription with *MYC* overexpression is suppressed upon KD of *CHD1*, returning to levels comparable to PE control cells (Fig. 6D-E). The overall reduction in EU signal is evident both in the nucleoplasm and in the nucleolus, where it represents rRNA synthesis (Fig. 6C). Taken together, these results indicate that CHD1 is required for the elevated levels of nascent transcription triggered by *MYC* dysregulation in human breast epithelial cells. In this setting, KD of *CHD1* leads to reduced transcriptional output and signs of nucleolar stress, potentially contributing to synthetic lethality with *MYC* dysregulation and suppression of tumor growth.

## DISCUSSION

In this study, we identify the chromatin remodeler CHD1 as a key mediator of MYC-driven tumorigenesis in a model of breast cancer. We show that CHD1 maintains an open chromatin landscape and a transcriptional program associated with transformation. *CHD1* knockdown reduces global levels of transcription concomitant with compromised nucleolar function, impairs survival and proliferation specifically upon MYC overexpression, ultimately impairing tumorigenesis (Fig. 6E). Taken together, our results provide new insights into the chromatin-level underpinnings of MYC-driven cancer and open several avenues of future inquiry.

MYC and CHD1 have several points in common at the functional level. They both bind to transcriptionally active open chromatin, marked by the activating histone mark H3K4me3 and high levels of RNA Pol I and II^4,21,22,44^. The double chromodomains of CHD1 bind directly and specifically to H3K4me2/3, and CHD1 and MYC have been detected as interacting proteins^45^. Additionally, CHD1 interacts with TOP2β, while MYC forms a complex with both TOP1 and TOP2 known as the topoisome^23,46,47^. Both CHD1 and MYC have independently been shown to promote transcriptional output in various cell types^4,22,26,27,44,48^. This high degree of overlap in genomic localization and function, together with the epistatic interaction revealed by this study, suggest that CHD1 acts downstream of MYC in maintain elevated RNA Pol I and II transcription. This may occur via CHD1 removing the +1 nucleosomal barrier to transcription and/or via promoting repair of transient DNA breaks induced by high rates of transcription^23,24^. It will be of interest to further dissect how the MYC/CHD1 relationship impacts the genomic location and activity of RNA Polymerases.

It remains unclear how *CHD1* KD impairs cell cycle progression and induces cell death in MYC-overexpressing cells. One possibility is that cells are unable to repair DNA breaks created by MYC-induced increases in transcriptional output when *CHD1* is depleted^23^. Another non-mutually exclusive possibility is defective transcription of ribosomal RNA triggering nucleolar stress and p53-mediated apoptosis.

Our results raise the possibility of pharmacologically targeting CHD1 in MYC-driven cancers. CHD1 is a large, ∼200kDa protein with three distinct domains: a double chromodomain that recognizes H3K4me2/3, a DNA helicase/ATPase domain, and a DNA binding domain^49^. Thus, there are ample opportunities to molecularly target CHD1 by small molecules. Interestingly, a recent report has put forth a CHD1 antagonist, UNC10142, which shows anti-tumor efficacy in *PTEN-*deficient prostate cancer^50^. This small molecule targets the double chromodomain of CHD1 in the H3K4me2/3-binding pocket, decreases binding of the protein to this histone mark in vitro and reduces viability of *PTEN-*deficient prostate cancer cells, with minimal impact in *PTEN-*intact cells^50^. These findings provide a proof-of-concept for the feasibility of developing small molecule inhibitors of CHD1, either by increasing the potency of UNC10142 or by targeting other functional regions of the protein. Interestingly, CHD1 has been reported to promote the proliferation of estrogen receptor-positive (ER+) breast cancer cells in response to estrogen, a context where MYC is required^51,52^. As the MCF10A lines used in this study are ER-, this suggests that targeting CHD1 may be broadly relevant in different subtypes of breast cancer. In addition to the requirement for CHD1 in MYC-driven breast cancer (this study) and in *PTEN*-deficient prostate cancer^53^, CHD1 has also been shown to be required for *KRAS*-driven anchorage-independent growth of human colon cancer cells^54^. In parallel, CHD1 was identified as a positive regulator of HIV transcription in an unbiased insertional mutagenesis screen^55^. Thus, CHD1 may be an important nexus for the regulation of transcriptional output in various physiological and pathological contexts.

## MATERIALS AND METHODS

### Cell culture

MCF10A PE and PM cells were cultured in DMEM/F12 (Invitrogen #111330-032) supplemented with 5% horse serum (Invitrogen #16050-122), 20ng/mL hEGF (Peprotech #AF-100-15), 500ng/mL hydrocortisone (Sigma #H-0888), 100ng/mL cholera toxin (Sigma #C8052-2MG), 10μg/mL insulin (Sigma #I-1882), and 100U/ml penicillin/streptomycin (Life Technologies #15140122). Cell lines were cultured at 37°C in a humidified incubator with 5% CO_2_ and 95% air. Dox-treatment was performed at a concentration of 50ng/mL.

### Short-hairpin RNA design

The shRNA targeting sequences were designed using the Broad Institute Genetic Perturbation Platform (https://portals.broadinstitute.org/gpp/public/). Two independent shRNAs targeting *CHD1* were generated. Forward and reverse oligos were annealed and cloned into the Tet-pLKO-puro backbone (Addgene #21915). One negative shRNA control (shGFP) was generated targeting GFP. shCHD1 A targets exon 24 of the *CHD1* gene, while shCHD1 B targets exon 28.

*CHD1 A (Forward):*

5’-CCGGGCGGTTTATCAAGAGCTATAACTCGAGTTATAGCTCTTGATAAAC-CGCTTTTTG-3’

*CHD1 A (Reverse):*

5’-AATTCAAAAAGCGGTTTATCAAGAGCTATAACTCGAGTTATAGCTCTTG-ATAAACCGC-3’

*CHD1 B (Forward):*

5’-CCGGCGTGCAGACTACCTCATCAAACTCGAGTTTGATGAGGTAGTCTGC-ACGTTTTTG-3’

*CHD1 B (Reverse):*

5’-AATTCAAAAACGTGCAGACTACCTCATCAAACTCGAGTTTGATGAGGTA-GTCTGCACG-3’

### Lentivirus generation, transduction and cell line validation

HEK293T cells were transfected in 15cm dishes at approximately 70-80% confluency with lentiviral packaging plasmids *pMD2.G* and *psPAX2* and the lentiviral DNA construct containing the shRNA plasmid following the Lipofectamine 2000 protocol (Invitrogen #11668027). Cell culture medium was changed the following day and viral medium was harvested and filtered through a 0.45µm filter at 48h post-transfection. To generate shRNA-containing PE and PM lines, cells were transduced with lentivirus in 0.8mg/mL polybrene for 24h. Cell culture medium was changed at 24h post-infection and puromycin (1µg/ml) selection began at 48h post-infection. *CHD1* KD was validated at the transcriptomic and protein levels using qRT-PCR and Western blot, respectively.

### Mouse xenograft

A panel of PM cells containing either Dox-inducible shGFP (negative control), shTOP1 (positive control; a gift from Dr. Linda Penn), shCHD1 A or shCHD1 B were injected into female NOD-SCID mice at 6-7 weeks old. 1×10^6^ cells were subcutaneously injected into the flanks of each mouse in 100µL volumes of 50% Matrigel (Corning #354262). Immediately after injection, mice were provided 50mg/mL Dox-supplemented drinking water (Sigma-Aldrich #D9891) for the duration of the experiment, which was refreshed twice a week. The experimental endpoint was defined to be when the first time a tumor volume of >1000mm^3^ was achieved, which occurred at 62 days. Tumor volume was measured twice a week and tumor mass was measured at endpoint. Animal Use Protocols (AUP972.35) describing this experiment was approved and overseen by the UHN Animal Resources Centre in compliance with the Canadian Council on Animal Care guidelines.

### Proliferation curve

A panel of PE/PM cells were plated at a 2.5×10^4^ cells per condition and immediately treated with Dox. Cells were harvested at 24-hour intervals and live cells were counted with an automated cell counter (Bio-Rad #1450102) in triplicate with Trypan blue (Bio-Rad #1450021).

### Clonogenic assay

A panel of PE/PM cells were treated with Dox for 24h before being plated in triplicate in 6-well plates at a density of 1×10^3^ cells per well. Colonies were grown undisturbed for 6 days, after which cells were washed twice in cold 1x PBS and fixed in ice-cold (−20°C) methanol for 10min. Cells were then stained with 0.5% crystal violet solution for 10min and washed with dH_2_O and air-dried overnight.

### 3D Matrigel assay

Pre-chilled 8-chamber BD Falcon CultureSlides (#08-774-26) were coated in cold Matrigel (VWR #CA89050-192) and allowed to solidify at 37°C for 45min. Cells were harvested, resuspended, and counted in assay media (DMEM/F12 supplemented with 2% horse serum, 500ng/mL hydrocortisone, 100ng/mL cholera toxin, 10μg/mL insulin, and 100U/ml penicillin/streptomycin). 1×10^3^ cells per each condition were resuspended in 400µL assay media supplemented with Dox, 5ng/mL EGF and 2% Matrigel and plated onto Matrigel-coated chamber slides. Cell culture medium was refreshed every 3 days with supplemented assay medium for a period of 14 days, at which point the total number of colonies were counted.

### AnnexinV/PI flow cytometry

Cells were plated in 6-well plates at a density of 2.5×10^4^ cells/well and treated with Dox for 96h. After Dox-treatment, cell culture medium was collected from each well and adherent cells were harvested. Samples were stained with 2µg/mL PI and AnnexinV-APC (Molecular Probes #A35110) in AnnexinV buffer (Biolegend #422201) for 15min in the dark at room temperature. Single colour and a positive cell-death control were also included. Samples were analyzed by flow cytometry on a Beckman Coulter Gallios and analyzed by Kaluza.

### RNA-sequencing and data analysis

Total RNA was extracted from 2×10^5^ live cells after 72h of Dox-treatment with the RNeasy Mini Kit. Total isolated RNA was quantified by Qubit HS RNA Kit (Invitrogen #Q32852) and RNA quality was measured with the Fragment Analyzer HS RNA Kit (Agilent #DNF-472). The RNA-seq library was generated following the manufacturer’s instructions for the NEBNext Poly(A) mRNA Magnetic Isolation Module (NEB #E7490S) and the NEBNext Ultra II Directional RNA Library Prep Kit (NEB #E7760L). cDNA library quality was assessed by Fragment Analyzer NGS Kit (Agilent #DNF-474). Libraries were quantified by Qubit HS DNA kit (Invitrogen #Q32851) and pooled at equimolar concentration. Libraries were paired end sequenced in an Illumina NovaSeqX 10B flow cell at The Centre for Applied Genomics.

Library quality was assessed using FastQC and trimmed of adaptor sequences with Trim Galore! v0.4.0 (Babraham Bioinformatics). Alignment was then performed using hisat2 (v2.2.1) to the GRCh38 reference genome^56^. Gene counts were obtained using the featureCounts function on Rsubread (v1.22.3). Data were analysed using tidyverse (v1.3.0) and visualized using ggplot2 (v3.3.5). Genes were considered differentially expressed and defined as significant if the adjusted P value was < 0.05, and if they had an absolute FC>1.5. GSEA analyses were performed with the fGSEA package (v1.24.0) and gene set collections were downloaded from the Molecular Signatures Database (https://www.gsea-msigdb.org/gsea/msigdb/index.jsp). Data of replicates from shRNA A and B were pooled for downstream analyses.

### EdU cell cycle analysis

Cells were seeded at a density of 1×10^5^ cells/plate and treated with of Dox for 72h. At 72h post-Dox treatment, cells were refreshed with medium containing 10µM EdU and Dox and incubated for 1 hour. Staining was performed following the Click-iT^™^ EdU Alexa Fluor^™^ 647 Flow Cytometry Assay Kit manufacturer’s protocol (Molecular Probes #C10424). Cells were incubated with FxCycle Violet Stain (Molecular Probes #F10347) and RNaseA (Life Technologies #EN0531) for 30min before proceeding directly to flow cytometry using a Beckman Coulter Gallios. Data was analyzed by FlowJo (v10.10).

### EU incorporation assay and immunofluorescence

Cells were plated on #1.5, 18mm circular glass coverslips (Fisher Scientific #50948975) in 12-well cell culture plates and treated with Dox for 72h. For EU incorporation, cells were refreshed with medium containing 1mM EU and Dox for 45min. Coverslips were fixed for 15min with 4% paraformaldehyde (VWR # CAAAJ61899-AP), washed 1x in PBS and permeabilized for 15min with 0.5% Triton X-100 in PBS (Sigma Aldrich #9036-19-5). For EU experiments, cells were washed once in 3% BSA in PBS and the subsequent Click-iT reaction, washes and DNA staining were performed as per the Click-iT^™^ RNA Alexa Fluor^™^ 488 Imaging Kit manufacturer’s protocol (Invitrogen #C10329). For regular immunofluorescence, cells were blocked with 3% donkey serum (Sigma-Aldrich D9663), 1% BSA in PBS-T. Primary antibody staining for NPM1 (1:200; Proteintech #60096-1) or p53-AF647 (1:200; Santa Cruz #sc-47698-AF647) was done overnight at 4°C. Cells were washed three times in PBS-T and incubated with fluorescence-conjugated secondary antibody for 1h in the dark at room temperature. Coverslips were washed 3x in PBS-T and mounted on 1mm Superfrost Plus Microscope slides (Fisher# 1255015) with Immu-Mount (Epredia #9990412). Images were captured on an Andor BC43 Benchtop Confocal Microscope. Quantification was performed in ImageJ with Z-projections as average intensity. Nuclei were quantified using the ROI manager with background correction.

### ATAC-seq

ATAC-seq libraries were generated following the Omni-ATAC protocol on PM shGFP, PM shCHD1 A, and PM shCHD1 B cells after 72h of Dox-treatment^57^. 50,000 live cells were sorted and then centrifuged at 500xg for 5min at 4°C. Cell pellets were then resuspended in 50µL of cold resuspension buffer (RSB; 10 mMTris-HCl at pH 7.4, 10mM NaCl, 3 mM MgCl2) supplemented with 0.1% NP-40, 0.1% Tween-20, and 0.01% digitonin, to dissociate cell membrane. Samples were incubated on ice for 3min and then washed with 1mL of ATAC RSB with 0.1% NP-40 by inverting the tube. The nuclei were subsequently pelleted by centrifugation at 500xg for 10min at 4°C and then resuspended in 50µL of transposition mixture (25µL of 2x TD buffer, 2.5µL of transposase [100nM final], 16.5µL of 1x PBS, 0.5µL of 1% digitonin, 0.5µL of 10% Tween-20, 5µL of nuclease-free water) for tagmentation. Tagmentation was performed at 37°C for 30min at 1000RPM on a thermomixer. DNA clean-up was performed following the tagmentation using the Zymo DNA Clean and Concentrator 5 kit (Zymo #D4014) and DNA was eluted in 21µL of elution buffer. Library amplification was performed as per the Omni-ATAC protocol^57^. After five initial amplification cycles, an additional qPCR surveillance step was included to determine the optimal number of extra PCR amplification cycles, enough to generate sufficient material for library sequencing, without over-amplifying and introducing duplicate artifacts. Amplified libraries were then purified with AMPure XP beads (Beckman Coulter #A63880) using a double-sided bead purification approach, to remove smaller primer dimers as well as large >1000bp fragments. Finally, libraries were quantified using the Qubit dsDNA high-sensitivity kit (Invitrogen #Q32854) and pooled at equimolar concentrations to be sequenced paired-end on a single lane of an Illumina NovaSeqX 10B flow cell.

Read quality was assessed using FastQC (v0.12.1), and results summarized with MultiQC (v1.25.2). Pair-end reads were trimmed using Trim Galore (v0.6.6). Reads that passed the quality control and trimming steps were aligned to GRCh38 using Bowtie2 (v2.4.1). Unaligned reads, mitochondrial reads, and duplicate reads were removed at this stage. Peak calling was performed with Macs3 (v3.0.3), using –nomodel and –nolambda parameters. Peaks overlapping with human blacklist regions were filtered out. As further validation steps, library was checked for Fraction of Reads in Peaks (FriP), enrichment of reads at promoters, fragment length distribution, and genome distribution. Differential accessibility analysis was performed with R (v4.4.1) using the DiffBind package (v3.16.0). For transcription factor motifs enrichment analysis, we first used the gkmSVM pipeline (v0.83.0) to select high-scoring kmers enriched in open and closed regions. These kmers were clustered using Starcode (v1.4). Each cluster was aligned using Clustal Omega (v1.2.4) and the resulting alignments were used to generate consensus motifs using MEME (v5.3.2). Motif logos and position weight matrices (PWMs) were produced. To identify candidate transcription factors binding these motifs, the generated PWMs were compared to known motif databases using TOMTOM (v5.5.7, online version).

## Competing interest statement

The authors declare no competing interests.

## Acknowledgments

We thank members of the Santos lab and J. Fish for critical reading of the manuscript. We are grateful to all members of the Santos and Penn labs for their input throughout the project. We thank A. Bang at the LTRI Flow Cytometry Facility, R. Bielecki and L. Brown at the LTRI Microscopy Facility, and S. Pereira at the SickKids TCAG Sequencing Facility. B.C. was supported by the Bank of Montreal Studentship in Medical Research at the Lunenfeld-Tanenbaum Research Institute. G.F. was supported by the Banting Postdoctoral Fellowship. This work was supported by CIHR Project Grant 156167 to L.Z.P. and Canada 150 Research Chair in Developmental Epigenetics and CIHR Project Grant 526948 to M.R.-S.

## Author contributions

M.R.-S, L.Z.P. and B.C. conceived the project and designed the experiments. B.C. performed most of the experiments and interpreted the data. G.F. contributed to analysis of RNA-seq data, ATAC-seq library preparation and analysis of ATAC-seq data. P.L. and M.S. performed and analyzed xenograft experiments. K.M. assisted with immunofluorescence. A.M. assisted with AnnexinV/PI analysis. M.R.-S. and L.Z.P. supervised the project. B.C. and M.R.-S. wrote the manuscript with input from all authors.

## Data and code availability

Sequencing data, including raw reads and processed data (raw counts, normalized counts, and differential gene expression table have been deposited on the NCBI Gene Expression Omnibus (GEO) repository (https://ncbi.nlm.nih.gov/geo) and will be accessible upon publication.

## Supplementary Figures

**Supplemental Figure S1.**
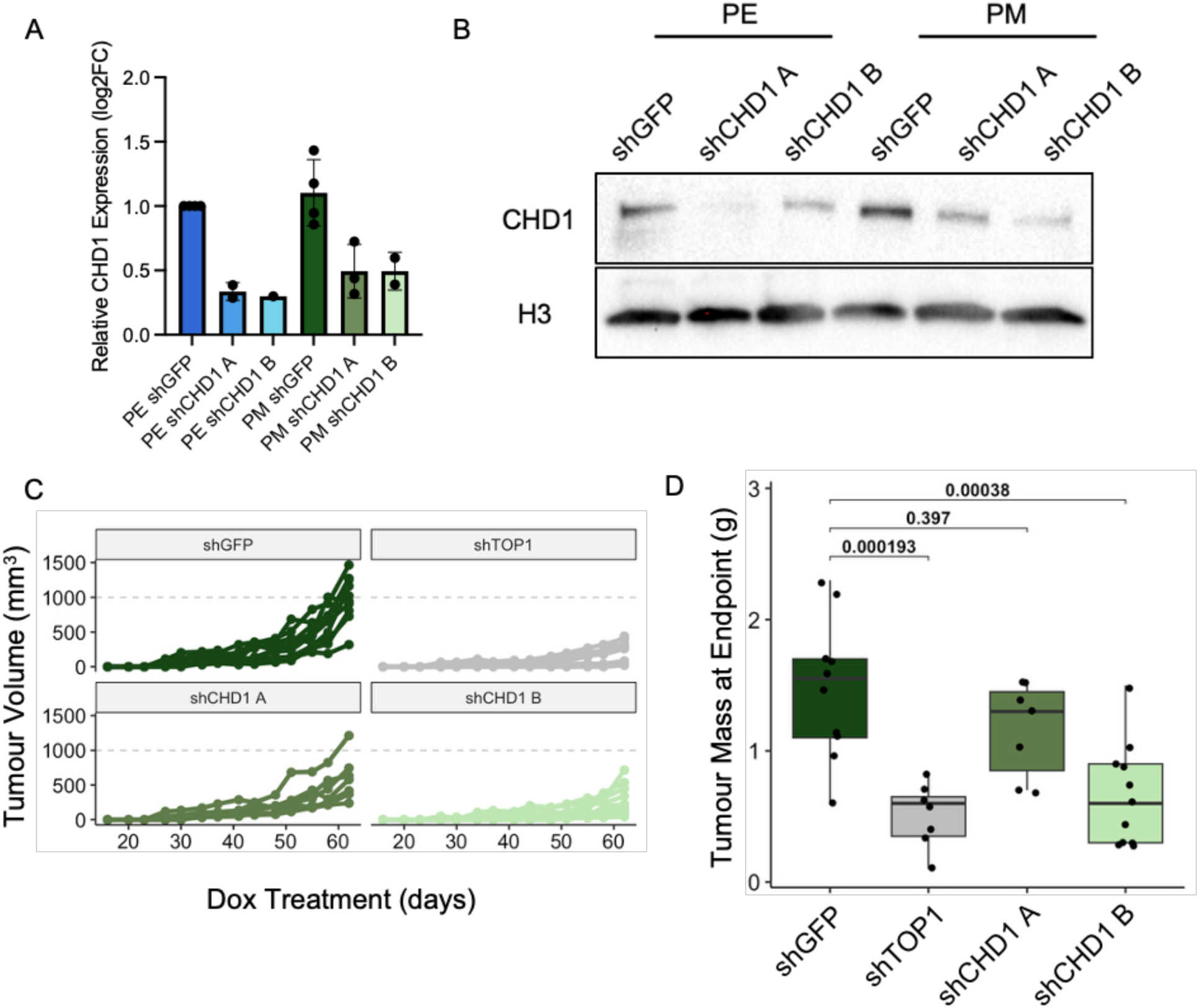
Validation of *CHD1* knockdown and additional xenograft data. A) qRT-PCR validation of *CHD1* knockdown following 48h of Dox treatment. Data are average fold change ± SEM, n = 2-4 biological replicates. B) Western blot probing for CHD1 protein with H3 as a loading control, n = 3 biological replicates. C) Tumor volume (mm^3^) in each individual mouse measured over the course of a xenograft experiment, n = 7-11 mice per condition. D) Tumor mass at endpoint in grams. P-values were calculated by one-way ANOVA with Tukey’s HSD.

**Supplemental Figure S2.**
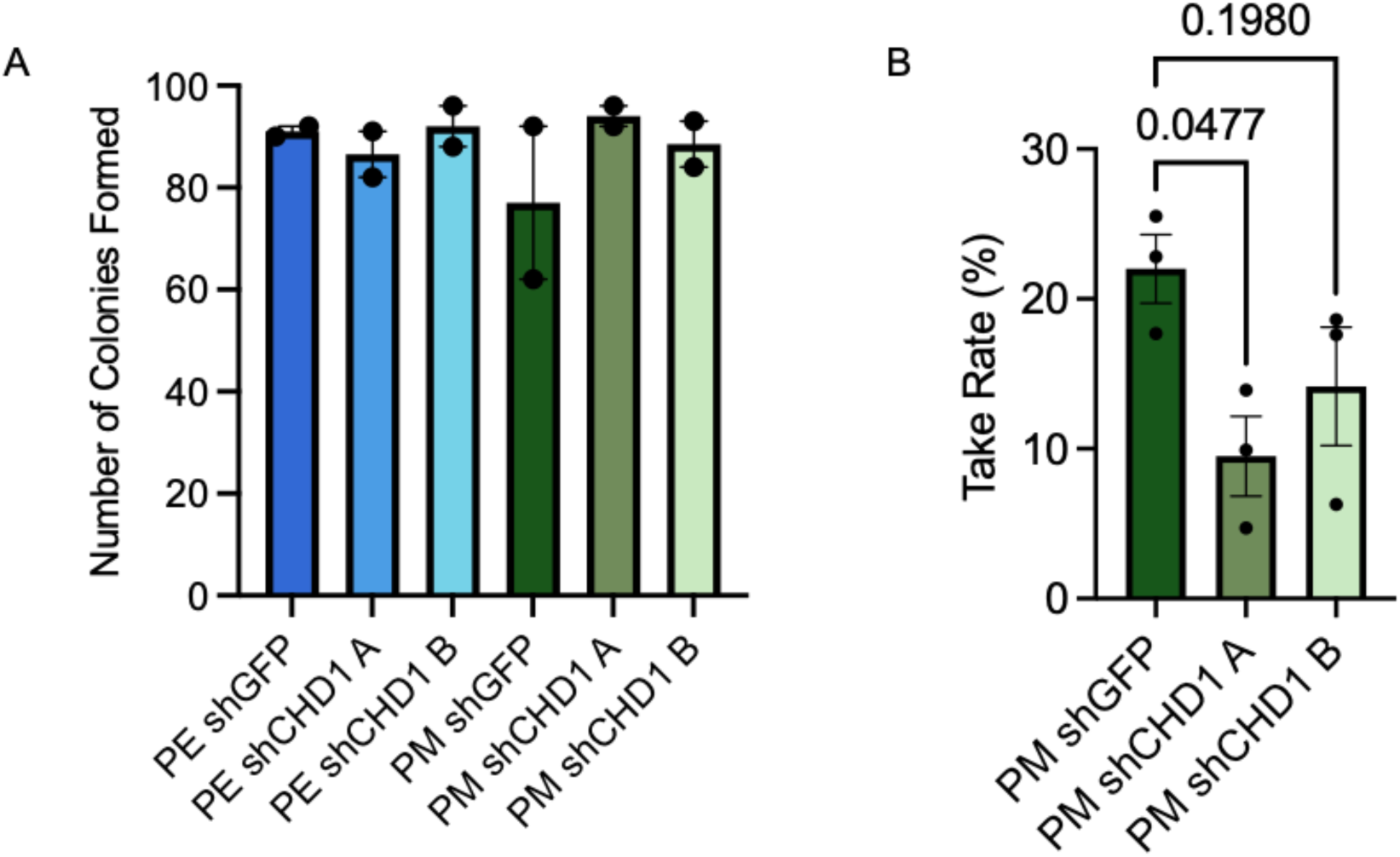
Dox-dependence and take rate in 3D cultures of *CHD1* KD cells. A) Number of colonies formed in a clonogenic assay using the 10A panel without doxycycline treatment, n=2 biological replicates per condition. Data is presented as mean ± SEM. B) Quantification of the take rate of cells by 3D Matrigel growth assays after 14 days of tamoxifen treatment. Take rate is defined as a percentage of the total organoids counted at endpoint over the total number of cells seeded at the start of the experiment. Data is presented as mean ± SEM. n=3 independent biological replicates. P-values were calculated by one-way ANOVA with Tukey’s multiple comparisons test.

**Supplemental Figure S3.**
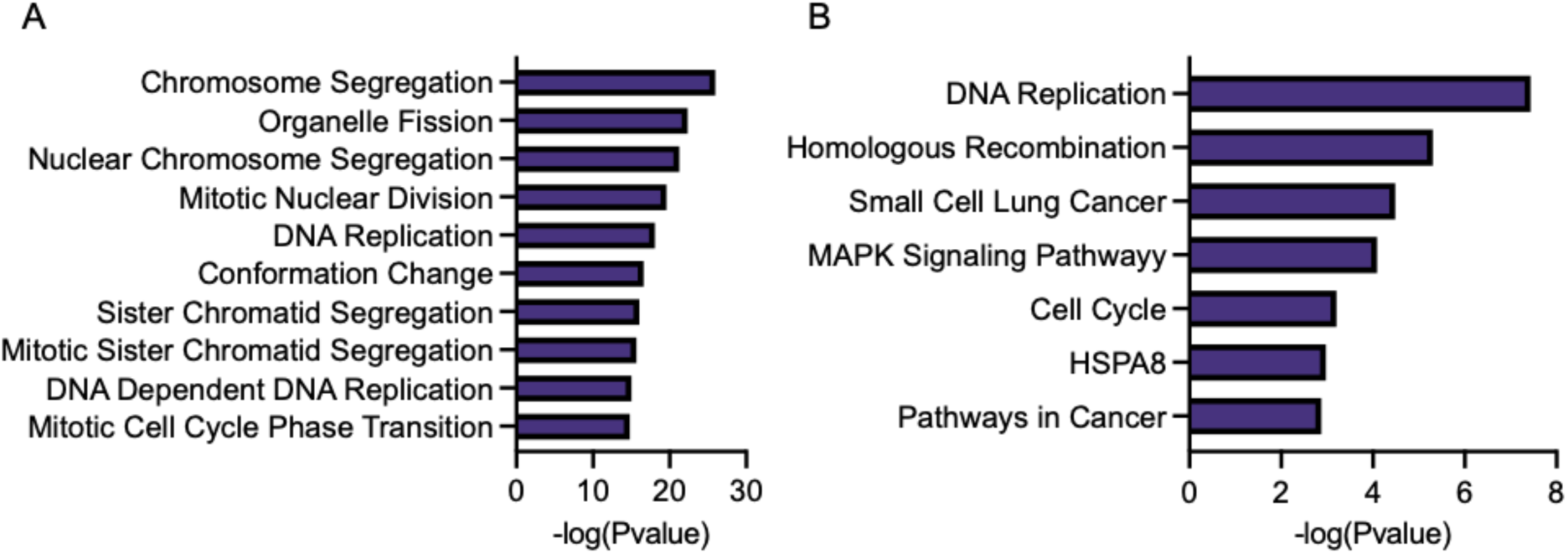
Functional terms enriched in downregulated genes upon *CHD1* KD. A) GO analysis of genes differentially expressed in both 10A.PM shCHD1 conditions relative to the PM shGFP controls. Shown are the top downregulated GO BP pathways ranked by –log(Pvalue). B) GO analysis of genes differentially expressed in both 10A.PM shCHD1 conditions relative to the PM shGFP controls. Shown are the most downregulated KEGG pathways ranked by –log(Pvalue).

**Supplemental Figure S4.**
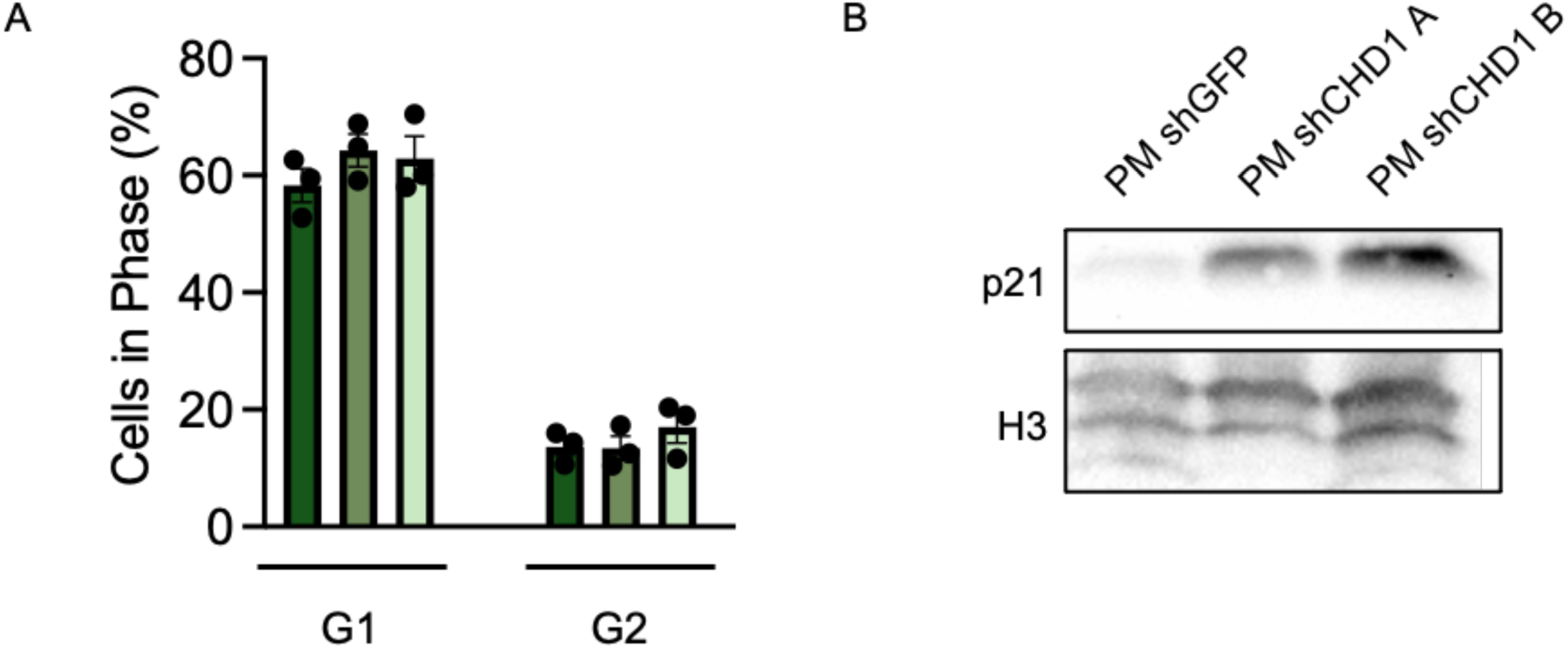
Supporting cell cycle data in *CHD1* KD cells. A) Quantification of the proportion of cells in G1 and G2 from EdU cell cycle analysis by flow cytometry (n=3 biological replicates). Data is presented as the mean ± SEM. B) Western blot analysis for p21 protein abundance in the panel of PM breast cancer lines. H3 is included as a loading control (n=2 biological replicates).

**Supplemental Figure 5.**
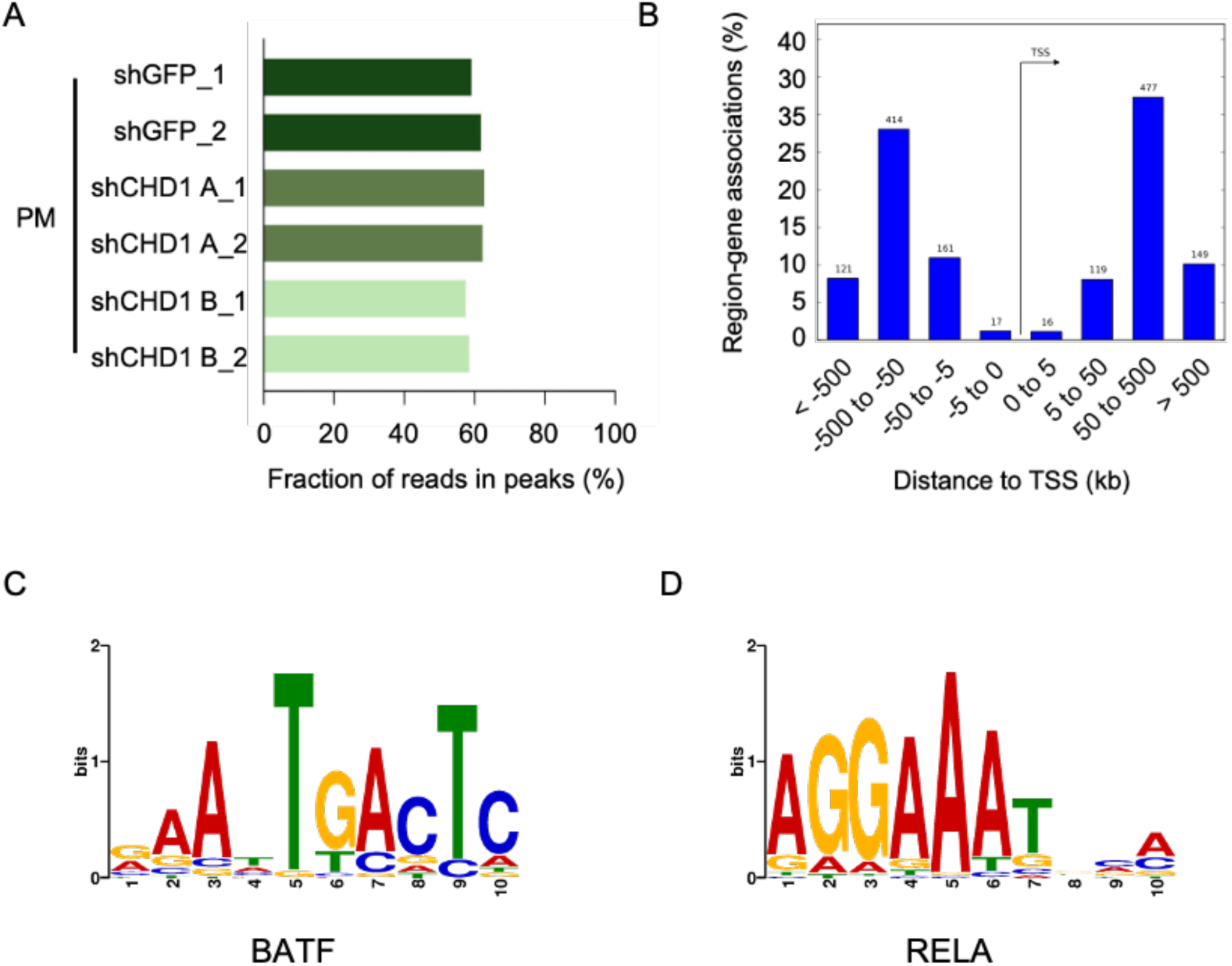
ATAC-seq validation and motifs enriched in regions with losses in chromatin accessibility upon *CHD1* KD. A) Fraction of reads in peaks for ATAC-seq samples. B) Genomic distribution of reads based on distance from TSS in ATAC-seq samples. C) DNA motif highly enriched in regions with a loss of chromatin accessibility upon *CHD1* KD, with the transcription factor that best matches the motif: BATF. D) DNA motif highly enriched in regions with a loss of chromatin accessibility upon *CHD1* KD, with the transcription factor that best matches the motif: RELA.

## Notes

### Competing Interest Statement

The authors have declared no competing interest.

## References

1. Dang, C. V. c-Myc Target Genes Involved in Cell Growth, Apoptosis, and Metabolism. Mol Cell Biol 19, 1–11 (1999).

2. Percharde, M., Wong, P. & Ramalho-Santos, M. Global Hypertranscription in the Mouse Embryonic Germline. Cell Rep 19, 1987–1996 (2017).

3. Collignon, E. et al. m6A RNA methylation orchestrates transcriptional dormancy during paused pluripotency. Nat Cell Biol 25, 1279–1289 (2023).

4. Nie, Z. et al. c-Myc Is a Universal Amplifier of Expressed Genes in Lymphocytes and Embryonic Stem Cells. Cell 151, 68–79 (2012).

5. Varlakhanova, N. V. et al. myc maintains embryonic stem cell pluripotency and self-renewal. Differentiation 80, 9–19 (2010).

6. Wilson, A. et al. c-Myc controls the balance between hematopoietic stem cell self-renewal and differentiation. Genes Dev 18, 2747–2763 (2004).

7. Lourenco, C. et al. MYC protein interactors in gene transcription and cancer. Nat Rev Cancer 21, 579–591 (2021).

8. Dong, Y., Tu, R., Liu, H. & Qing, G. Regulation of cancer cell metabolism: oncogenic MYC in the driver’s seat. Signal Transduct Target Ther 5, 124 (2020).

9. Kalkat, M. et al. MYC deregulation in primary human cancers. Genes (Basel) 8, 2–30 (2017).

10. Duffy, M. J., O’Grady, S., Tang, M. & Crown, J. MYC as a target for cancer treatment. Cancer Treat Rev 94, 102154 (2021).

11. Madden, S. K., de Araujo, A. D., Gerhardt, M., Fairlie, D. P. & Mason, J. M. Taking the Myc out of cancer: toward therapeutic strategies to directly inhibit c-Myc. Mol Cancer 20, 3 (2021).

12. Thng, D. K. H., Toh, T. B. & Chow, E. K. H. Capitalizing on Synthetic Lethality of MYC to Treat Cancer in the Digital Age. Trends Pharmacol Sci 42, 166–182 (2021).

13. Kessler, J. D. et al. A SUMOylation-dependent transcriptional subprogram is required for Myc-driven tumorigenesis. Science 335, 348–53 (2012).

14. Huang, C. H. et al. CDK9-mediated transcription elongation is required for MYC addiction in hepatocellular carcinoma. Genes Dev 28, 1800–1814 (2014).

15. Toyoshima, M. et al. Functional genomics identifies therapeutic targets for MYC-driven cancer. Proc Natl Acad Sci U S A 109, 9545–9550 (2012).

16. Horiuchi, D. et al. MYC pathway activation in triple-negative breast cancer is synthetic lethal with CDK inhibition. J Exp Med 209, 679–696 (2012).

17. Einstein, J. M. et al. Inhibition of YTHDF2 triggers proteotoxic cell death in MYC-driven breast cancer. Mol Cell 81, 3048–3064.e9 (2021).

18. Sato, M. et al. The UVSSA complex alleviates MYC-driven transcription stress. J Cell Biol 220, (2021).

19. Kessler, J. D. et al. A SUMOylation-Dependent Transcriptional Subprogram Is Required for Myc-Driven Tumorigenesis. Science (1979) 335, 348–353 (2012).

20. Lin, P. et al. Topoisomerase 1 Inhibition in MYC-Driven Cancer Promotes Aberrant R-Loop Accumulation to Induce Synthetic Lethality. Cancer Res 183, 4015–4029 (2023).

21. Gaspar-Maia, A. et al. Chd1 regulates open chromatin and pluripotency of embryonic stem cells. Nature 460, 863–868 (2009).

22. Guzman-Ayala, M. et al. Chd1 is essential for the high transcriptional output and rapid growth of the mouse epiblast. Development 142, 118–127 (2015).

23. Bulut-Karslioglu, A. et al. Chd1 protects genome integrity at promoters to sustain hypertranscription in embryonic stem cells. Nat Commun 12, 1–11 (2021).

24. Skene, P. J., Hernandez, A. E., Groudine, M. & Henikoff, S. The nucleosomal barrier to promoter escape by RNA polymerase II is overcome by the chromatin remodeler Chd1. Elife 3, 1–19 (2014).

25. Kim, Y. K. et al. Absolute scaling of single-cell transcriptomes identifies pervasive hypertranscription in adult stem and progenitor cells. Cell Rep 42, 111978 (2023).

26. Percharde, M., Bulut-Karslioglu, A. & Ramalho-Santos, M. Hypertranscription in Development, Stem Cells, and Regeneration. Dev Cell 40, 9–21 (2017).

27. Koh, F. M. et al. Emergence of hematopoietic stem and progenitor cells involves a Chd1-dependent increase in total nascent transcription. Proceedings of the National Academy of Sciences 112, E1734–E1743 (2015).

28. Bulut-Karslioglu, A. et al. The Transcriptionally Permissive Chromatin State of Embryonic Stem Cells Is Acutely Tuned to Translational Output. Cell Stem Cell 22, 369–383.e8 (2018).

29. Lourenco, C. et al. Modelling the MYC-driven normal-to-tumour switch in breast cancer. Dis Model Mech 12, 1–9 (2019).

30. Hanahan, D. & Weinberg, R. A. Hallmarks of cancer: The next generation. Cell 144, 646–674 (2011).

31. Liberti, M. V & Locasale, J. W. The Warburg Effect: How Does it Benefit Cancer Cells? Trends Biochem Sci 41, 211–218 (2016).

32. Zhou, C., Gao, Y., Ding, P., Wu, T. & Ji, G. The role of CXCL family members in different diseases. Cell Death Discov 9, 212 (2023).

33. Martínez-Pérez, C. et al. The Signal Transducer IL6ST (gp130) as a Predictive and Prognostic Biomarker in Breast Cancer. J Pers Med 11, 618 (2021).

34. Murakami, M., Kamimura, D. & Hirano, T. Pleiotropy and Specificity: Insights from the Interleukin 6 Family of Cytokines. Immunity 50, 812–831 (2019).

35. Vega-Benedetti, A. F. et al. PLAGL1 gene function during hepatoma cells proliferation. Oncotarget 9, 32775–32794 (2018).

36. Clouaire, T. et al. The THAP domain of THAP1 is a large C2CH module with zinc-dependent sequence-specific DNA-binding activity. Proc Natl Acad Sci U S A 102, 6907–6912 (2005).

37. Roussigne, M., Cayrol, C., Clouaire, T., Amalric, F. & Girard, J. P. THAP1 is a nuclear proapoptotic factor that links prostate-apoptosis-response-4 (Par-4) to PML nuclear bodies. Oncogene 22, 2432–2442 (2003).

38. Ataei, L. et al. LINE1 elements at distal junctions of rDNA repeats regulate nucleolar organization in human embryonic stem cells. Genes Dev 39, 280–298 (2025).

39. Yang, K., Yang, J. & Yi, J. Nucleolar stress: Hallmarks, sensing mechanism and diseases. Cell Stress 2, 125–140 (2018).

40. Zhang, J. et al. LINE1 and PRC2 control nucleolar organization and repression of the 8C state in human ESCs. Dev Cell 60, 186–203 (2025).

41. Potapova, T. A. et al. Distinct states of nucleolar stress induced by anticancer drugs. Elife 12, RP88799 (2023).

42. Xie, S. Q. et al. Nucleolar-based Dux repression is essential for embryonic two-cell stage exit. Genes Dev 34, 331–347 (2022).

43. Noller, H. F. The ribosome comes to life. Cell 187, 6486–6500 (2024).

44. Lin, C. Y. et al. Transcriptional amplification in tumor cells with elevated c-Myc. Cell 151, 56–67 (2012).

45. Heidelberger, J. B. et al. Proteomic profiling of VCP substrates links VCP to K6-linked ubiquitylation and c-Myc function. EMBO Rep 19, 1–20 (2018).

46. Das, S. K. et al. Excessive MYC-topoisome activity triggers acute DNA damage, MYC degradation, and replacement by a p53-topoisome. Mol Cell 84, 4059-4078.e10 (2024).

47. Das, S. K. et al. MYC assembles and stimulates topoisomerases 1 and 2 in a ‘topoisome’. Mol Cell 82, 140–158.e12 (2022).

48. Bulut-Karslioglu, A. et al. The Transcriptionally Permissive Chromatin State of Embryonic Stem Cells Is Acutely Tuned to Translational Output. Cell Stem Cell 22, 369–383.e8 (2018).

49. Stokes, D. G. & Perry, R. P. DNA-Binding and Chromatin Localization Properties of CHD1. Mol Cell Biol 15, 2745–2753 (1995).

50. Johnson, R. L. et al. Discovery of CHD1 Antagonists for PTEN-Deficient Prostate Cancer. J Med Chem 67, 20056–20075 (2024).

51. Tan, S. et al. Identification of miR-26 as a key mediator of estrogen stimulated cell proliferation by targeting CHD1, GREB1 and KPNA2. Breast Cancer Research 16, 1–13 (2014).

52. Cheng, A. S. L. et al. Combinatorial Analysis of Transcription Factor Partners Reveals Recruitment of c-MYC to Estrogen Receptor-α Responsive Promoters. Mol Cell 21, 393–404 (2006).

53. Zhao, D. et al. Synthetic essentiality of chromatin remodelling factor CHD1 in PTEN-deficient cancer. Nature 542, 484–488 (2017).

54. Yu, B. et al. Oncogenesis driven by the Ras/Raf pathway requires the SUMO E2 ligase Ubc9. Proc Natl Acad Sci U S A 112, E1724–33 (2015).

55. Rodgers, M. J., Banks, D. J., Bradley, K. A. & Young, J. A. CHD1 and CHD2 are positive regulators of HIV-1 gene expression. Virol J 11, 180 (2014).

56. Kim, D., Paggi, J. M., Park, C., Bennett, C. & Salzberg, S. L. Graph-based genome alignment and genotyping with HISAT2 and HISAT-genotype. Nat Biotechnol 37, 907–915 (2019).

57. Corces, M. R. et al. An improved ATAC-seq protocol reduces background and enables interrogation of frozen tissues. Nat Methods 14, 959–962 (2017).

